# Bacterial endosymbionts enhance fungal virulence by disrupting the disease-suppressive rhizobiome

**DOI:** 10.64898/2026.03.19.712832

**Authors:** Xingang Zhou, Xianhong Zhang, Lingyi Ran, Dongli Liu, Huiting Jia, Jingyu Zhang, Nan Zhang, Muhammad Khashi u Rahman, Alexandre Jousset, Francisco Dini-Andreote, Fengzhi Wu, Zhong Wei

## Abstract

The mechanisms by which bacterial endosymbionts (bacteria living within fungal cells) enhance the fitness and virulence of fungal pathogens remain poorly understood. Here, we report that the tomato Fusarium wilt pathogen *Fusarium oxysporum* f. sp. *lycopersici* (FOL) hosts *Achromobacter* sp. endosymbionts that enhance fungal virulence. This virulence potentiation is partially dependent on interactions with the native rhizosphere microbiota. We show that bacterial endosymbiont-harboring FOL reshapes the rhizosphere bacterial community during pathogen infection and decreased the abundance of disease□suppressive bacteria, including *Streptomyces* sp. taxa. This inhibitory effect is mediated by bacterial endosymbiont□stimulated production of beauvericin (an antibacterial cyclic hexadepsipeptide) by FOL. Together, our findings reveal a tripartite interaction in which a fungal pathogen leverages its bacterial endosymbiont to weaken rhizosphere microbiome□based disease suppression by inhibiting plant-protective bacterial taxa. This work highlights how cross□kingdom symbioses can modulate pathogen ecology and virulence in soil environments.

## INTRODUCTION

The majority of eukaryotes live in close association with bacterial endosymbionts.^1,2^ Interactions between hosts and distinct bacterial taxa can confer a wide range of fitness benefits, including modulation of host metabolism that enhances growth, reproduction, and tolerance to biotic and abiotic stressors.^1–5^ Among microbial eukaryotes, bacterial endosymbionts are found in close association with diverse fungal lineages, including saprotrophs, symbionts, and pathogens.^6,7^ Fungal phytopathogens, in particular, pose major threats to global food security and agricultural sustainability.^8^ Collectively, phytopathogens are responsible for approximately 14% of global crop yield losses and account for 17.5% of all pesticide applications worldwide.^9,10^ Despite extensive characterization of the molecular mechanisms mediating fungal pathogenicity in diverse crop systems (e.g., effector secretion, toxin production, and enzymatic degradation of plant cell walls),^11^ many aspects of fungal virulence and host resistance remain poorly understood. Specifically, recent studies have shown that bacterial endosymbionts of phytopathogenic fungi can play active roles in pathogenicity by enhancing resistance to natural enemies, suppressing plant immune responses, and increasing fungal virulence^1,6,7,12–14^. In several soil-borne phytopathogens, specific bacterial endosymbionts are essential for full virulence, including those associated with *Rhizopus microsporus, Fusarium fujikuroi*, and *Rhizoctonia solani*.^14–16^

In addition to the innate immune defenses, plants rely on their associated microbiota as an important line of defense against pathogen invasion.^2,17,18^ These so-called disease-suppressive plant-associated microorganisms can inhibit pathogen growth, attenuate virulence, and/or induce systemic resistance in plants.^2,18,19^ Increasing evidence supports that specific microbial taxa function as integral components of the plant immune system. That is, upon pathogen detection or infection, plants can actively recruit disease-suppressive bacteria, thereby strengthening immune responses.^20–23^ However, this microbial line of defense is not invulnerable. Fungal pathogens have evolved counterstrategies, including the secretion of virulence factors with antimicrobial activity, acting on the suppression of plant-beneficial bacteria during root infection.^24–27^ Consequently, breaching these microbial protective barriers may be as critical for successful infection as overcoming the plant’s innate immune defenses.^24^

Here, we investigated bacterial endosymbionts associated with the soil-borne fungal pathogen *Fusarium oxysporum* f. sp. *lycopersici* (FOL), the causal agent of Fusarium wilt of tomato (*Solanum lycopersicum*). *F. oxysporum* is among the top ten most destructive phytopathogenic fungi worldwide and is responsible for vascular wilt diseases in approximately 100 plant species.^28^ Specifically, this study aimed to determine whether bacterial endosymbionts associated with FOL contribute to pathogen infection by altering microbial-mediated disease-suppressive barriers within the plant rhizosphere. Members of the genus *Fusarium* are frequently associated with distinct bacterial taxa residing within fungal hyphae,^16,29^ suggesting potential functional roles for these bacterial endosymbionts in fungal biology and virulence. Building on this observation, we compared the infection performance of FOL in association with bacterial endosymbiont and FOL endosymbiont-free lines with respect to their effects on plant infection and shifts in the rhizosphere microbiota. We hypothesized that specific bacterial endosymbionts (*1*) enhance FOL virulence, and (*2*) modulate the effects of FOL infection on the structure of the resident rhizosphere microbiota.

## RESULTS

### Detection and taxonomic information of bacterial endosymbionts of FOL

To assess the presence of bacterial endosymbionts in FOL, we performed diagnostic PCR targeting the bacterial 16S ribosomal RNA (rRNA) gene. PCR amplification of FOL genomic DNA using the bacteria-specific primer pair 27F/1492R^30^ resulted in positive amplification fragments with approximately 1,400 bp (Figure 1A), consistent with the expected size of the bacterial 16S rRNA gene. To independently confirm the intracellular localization and viability of bacterial cells, Live/Dead staining was performed, followed by fluorescence microscopy. In brief, the obtained images revealed the presence of green-fluorescent, bacteria-like structures within FOL hyphae (Figure 1B), indicating the presence of viable bacterial endosymbionts in the fungal mycelium.

We performed a bacterial 16S rRNA gene clone library using the amplified PCR fragments to characterize the associated bacteria. Analysis of 29 clones revealed three distinct 16S rRNA gene sequences based on 100% nucleotide identity, with Clone sequence-5 as the dominant taxon (representing 26 of the 29 cloned sequences). All cloned sequences were taxonomically assigned to *Achromobacter xylosoxidans* (Figure S1A). To validate these findings and recover culturable endosymbionts, 23 bacterial colonies were isolated from mechanically disrupted hyphae of surface-sterilized FOL. These obtained bacterial isolates exhibited 100% 16S rRNA gene sequence identity to the Clone sequence-5 (Figure S1A). In addition, when grown on nutrient agar, these isolates formed circular, smooth, and translucent colonies (Figure 1C). One isolate, designated EB05, was selected as a representative strain for subsequent experiments. Transmission electron microscopy revealed that EB05 cells exhibited rod- to oval-shaped morphology (Figure 1C).

### Contribution of bacterial endosymbionts to FOL virulence

We evaluated the potential contribution of bacterial endosymbionts to the virulence of FOL. First, we assessed whether the bacterial endosymbiont EB05 alone exhibited pathogenicity to tomato plants. Following inoculation into sterile soil, EB05 was detected in the rhizosphere and root tissues of tomato plants (Figure S1B). However, EB05 did not induce visible disease symptoms in roots, stems, or leaves, nor did it significantly affect plant dry biomass (Figure S1C and S1D). Second, to directly test the role of bacterial endosymbionts in fungal virulence, a bacteria-free FOL strain (hereafter referred to as Cured-FOL) was generated by treating wild-type FOL with antibiotics minocycline and ciprofloxacin (Figure S1E). The bacterial endosymbiont EB05 was subsequently reintroduced into Cured-FOL by coculturing EB05 with the Cured-FOL, yielding a restored strain (Restored-FOL).^31^ Successful colonization of EB05 in Restored-FOL, and its absence in Cured-FOL, were confirmed by bacterial 16S rRNA gene detection and fluorescence microscopy (Figure 1A, 1B). Re-isolated of bacteria from Restored-FOL followed by 16S rRNA gene sequencing further verified successful recolonization by EB05. Vertical transmission of EB05 within the fungal host was confirmed, as EB05 was consistently isolated after nine successive subcultures of Restored-FOL (Figure 1A). Quantitative PCR analysis showed that bacterial colonization efficiency (expressed as the ratio of bacterial to fungal gene abundance), did not significantly differ between Restored-FOL and wild-type FOL (Figure S1F). Scanning electron microscopy revealed no detectable differences in hyphal morphology among wild-type FOL, Cured-FOL, and Restored-FOL (Figure S1E). Similarly, mycelial growth rates were comparable across all three strains. However, conidial production was higher in wild-type FOL compared with both Cured-FOL and Restored-FOL (Tukey’s HSD test, *P* = 1.44 × 10^−5^; Figure S1G).

To evaluate the contribution of bacterial endosymbionts to FOL virulence under different soil microbial contexts, we conducted a pot experiment using both natural soil (containing an intact soil microbiota) and sterile soil (microbiota-depleted). Results revealed that tomato plants inoculated with Cured-FOL consistently exhibited significantly lower disease severity and reduced FOL abundance in the rhizosphere compared with plants inoculated with bacterial endosymbiont-harboring strains (*i*.*e*., wild-type FOL and Restored-FOL) (Tukey’s HSD test, *P* = 7.74 × 10^−16^; Figure 1D and S1H). Notably, the magnitude of bacterial endosymbiont-mediated virulence (*i*.*e*., the difference in disease severity between bacterial endosymbiont-harboring and endosymbiont-free FOL strains) was higher in natural soil than in sterile soil (Tukey’s HSD test, *P* = 3.73 × 10^−7^; Figure 1E). In addition, the soil microbiota-mediated disease suppression (*i*.*e*., the difference in disease severity between sterile and natural soils) was stronger for the Cured-FOL treatment than the wild-type FOL and Restored-FOL treatments (Tukey’s HSD test, *P* = 2.44 × 10^−7^; Figure 1F). Collectively, these results indicate that bacterial endosymbionts enhance FOL virulence and that the extent of virulence enhancement is modulated by interactions with the rhizosphere microbiota.

### Bacterial endosymbionts mediate the effects of FOL on tomato rhizosphere microbiota

We examined the effects of different FOL strains on the structure of tomato rhizosphere bacterial community using 16S rRNA gene amplicon sequencing. Relative to the non-inoculated control, inoculation with all FOL strains resulted in significant shifts in rhizosphere bacterial communities, as indicated by changes in β-diversity (permutational multivariate analysis of variance, PERMANOVA, *R*^2^ = 0.322, *P* = 0.030 for wild-type FOL; *R*^2^ = 0.251, *P* = 0.030 for Cured-FOL; *R*^2^ = 0.317, *P* = 0.029 for Restored-FOL; Figure 2A]. Importantly, the rhizosphere bacterial community of the Cured-FOL treatment differed from those of the wild-type FOL and Restored-FOL treatments (PERMANOVA, *R*^2^ = 0.295, *P* = 0.031 for Cured-FOL vs. wild-type FOL; *R*^2^ = 0.280, *P* = 0.028 for Cured-FOL vs. restored FOL; Figure 2A). Consistent with these patterns, bacterial community dissimilarity between the control and Cured-FOL treatments was lower than that between the control and either wild-type FOL or Restored-FOL treatments (Tukey’s HSD test, *P* =6.77 × 10^−11^).

For downstream analyses, bacterial operational taxonomic units (OTUs) from this experiment were designed as FOL-responsive OTUs (f.OTUs). Compared with the non-inoculated control, inoculation with the wild-type FOL, Cured-FOL, and Restored-FOL increased the relative abundance of 26, 23, and 26 f.OTUs, respectively, and decreased the relative abundance of 51, 22, and 45 f.OTUs, respectively (DESeq2 analysis with Benjamini-Hochberg correction, adjusted *P* < 0.05; Figure 2B and S2). These differentially abundant f.OTUs were mostly affiliated with the phyla Actinobacteria and Proteobacteria (Table S1). Notably, ten Actinobacteria f.OTUs [including five *Streptomyces* sp. f.OTUs (f.OTU273, f.OTU2295, f.OTU5011, f.OTU5685 and f.OTU6745)], exhibited lower relative abundances in the wild-type FOL and Restored-FOL treatments, but higher relative abundances in the Cured-FOL treatment (Figure 2C). Concordantly, both the relative abundance (based on 16S rRNA gene amplicon sequencing) and absolute abundance (based on quantitative PCR) of *Streptomyces* sp. decreased in the wild-type FOL and Restored-FOL treatments but increased in the Cured-FOL treatment (Tukey’s HSD test, *P* = 1.14 × 10^−9^ and *P* = 1.44 × 10^−7^, respectively; Figure 2D).

### Specific *Streptomyces* sp. taxa protect tomato plants against FOL

To investigate the potential function of *Streptomyces* taxa whose abundances were altered by bacterial endosymbiont-harboring FOL, a total of 225 bacterial isolates were obtained from tomato rhizosphere. After removal of clonal duplicates (*i*.*e*., isolates sharing 100% 16S rRNA gene sequence identity), 17 *Streptomyces* isolates were recovered (Figure 3A). Among these, one representative strain, *S. ciscaucasicus* St52, which exhibited 100% 16S rRNA gene sequence identity to f.OTU5685 (Figure 3A; Table S2), was selected for further characterization. *In vitro* assays conducted on potato dextrose agar demonstrated that St52 inhibited the mycelium growth of wild-type FOL (Figure 3A). Consistent with this antagonistic activity, pot experiments revealed that soil inoculation with St52 reduced Fusarium wilt disease severity (Welch’s *t*-test, *P* = 9.57 × 10^−10^, *P* = 5.93 × 10^−9^, and *P* = 5.78 × 10^−8^ for wild-type FOL, Cured-FOL, and Restored-FOL, respectively) and decreased FOL abundance in the rhizosphere (Welch’s *t*-test, *P* = 1.54 × 10^−7^, *P* = 1.53 × 10^−13^, and *P* = 5.46 × 10^−8^ for wild-type FOL, Cured-FOL, and Restored-FOL, respectively; Figure 3B). Notably, the magnitude of disease suppression conferred by St52 was greater in plants challenged with Cured-FOL (60.04%) than in those challenged with wild-type FOL (31.83%) and Restored-FOL (37.46%), indicating that bacterial endosymbionts within FOL partially attenuate the protective effects of antagonistic *Streptomyces* sp.

### Bacterial endosymbionts modulate the inhibitory effects of FOL on *Streptomyces* sp. St52

We examined the effects of FOL and its bacterial endosymbiont EB05 on the abundance of the plant-beneficial bacterium St52 in tomato rhizosphere. Tomato plants grown in sterile soil were first inoculated with St52, and after ten days, were challenged with either FOL or EB05. Compared with the St52-only control, inoculation with wild-type FOL and Restored-FOL decreased St52 abundance in the rhizosphere (Welch’s *t*-test, *P* = 2.44 × 10^−9^ and *P* = 2.33 × 10^−6^, respectively; Figure 3C). In contrast, inoculation with Cured-FOL increased St52 abundance in the rhizosphere (Welch’s *t*-test, *P* = 9.01 × 10^−9^). These patterns are consistent with the rhizosphere community-level responses observed in the 16S rRNA gene amplicon sequencing analysis (Figure 2). Notably, inoculation with EB05 did not significantly affect St52 abundance in the rhizosphere.

We further evaluated the impact of FOL and EB05 on St52 growth in soil microcosms and *in vitro* co-culture assays. In both systems, wild-type FOL and Restored-FOL reduced St52 abundance in the soil (Welch’s *t*-test, *P* = 6.87 × 10^−13^ and *P* = 8.50 × 10^−12^, respectively; Figure 3D) and in oatmeal broth (Welch’s *t*-test, *P* = 5.60 × 10^−14^ and *P* = 1.51 × 10^−11^, respectively; Figure 3E). In contrast, neither Cured-FOL nor EB05 alone affected St52 abundance under these conditions. Collectively, these results demonstrate that bacterial endosymbionts enhance the inhibitory effects of FOL on *Streptomyces* sp. ST52, whereas the bacterial endosymbiont EB05 alone is insufficient to suppress St52 growth.

### Bacterial endosymbionts enhance the production of beauvericin in FOL

We investigated whether bacterial endosymbionts modulate FOL metabolism in a manner that contributes to the inhibition of St52. To this end, we first evaluated the effects of the cell-free culture supernatants from FOL strains and EB05 on the *in vitro* growth of St52. Supernatants derived from wild-type FOL and Restored-FOL inhibited St52 growth (Welch’s *t*-test, *P* = 5.12 × 10^−10^ and *P* = 6.99 × 10^−9^, respectively; Figure 4A), whereas supernatants from Cured-FOL and EB05 had no inhibitory effect. Further high-performance liquid chromatography-mass spectrometry (HPLC-MS) analysis revealed that the chemical profiles of supernatants from wild-type FOL and Restored-FOL differed from those of Cured-FOL (Permutation test, *R*^*2*^*X* = 0.330 and *Q*^*2*^ = 0.680 for wild-type FOL vs. Cured-FOL, *R*^*2*^*X* = 0.342 and *Q*^*2*^ = 0.742 for Restored-FOL vs. Cured-FOL; Figure S3A and S3B). Eight compounds, *i*.*e*., beauvericin, N-acetylarylamine, methylglutaric acid, phenol glucuronide, N-acetyl-L-tyrosine, 2-phenylbutyric acid, 2-hydroxystearic acid, and ethyl tetradecanoate, were present at higher concentrations in supernatants from wild-type FOL and Restored-FOL compared with Cured-FOL (variable importance of projection value > 1.5, log_2_ fold change > 1 and Benjamini-Hochberg adjusted *P* < 0.01; Figure 4B). Subsequent *in vitro* assays demonstrated that among these compounds, only beauvericin (at concentrations of 0.1 − 10 μg ml^−1^) inhibited the growth of St52 (Tukey’s HSD test, *P* = 2.20 × 10^−16^; Figure 4C and S3C). In contrast, beauvericin at these concentrations did not affect the growth of EB05 (Figure S3D). When tested against three additional bacterial strains isolated from tomato rhizosphere (i.e., *Bacillus, Pseudomonas*, and *Sphingomonas* spp.), beauvericin inhibited bacterial growth only at the highest concentration tested (*i*.*e*., 10 μg ml^−1^) (Welch’s *t*-test, *P* = 4.51 × 10^−4^, *P* = 1.56 × 10^−4^ and *P* = 3.43 × 10^−5^, respectively; Figure S3D). Finally, beauvericin at 10 μg ml^−1^ induced reactive oxygen species (*i*.*e*., hydrogen peroxide) in tomato roots (Figure S3E), indicating phytotoxic activity.^32^

We further validated that the contribution of bacterial endosymbionts to FOL virulence is associated with beauvericin production. Beauvericin concentrations were quantified in the rhizosphere of tomato plants inoculated with FOL. HPLC analysis revealed that beauvericin levels in the rhizosphere of plants inoculated with Cured-FOL were lower than those in plants inoculated with wild-type FOL or Restored-FOL (Welch’s *t*-test, *P* = 1.86 × 10^−7^ and *P* = 1.86 × 10^−6^, respectively; Figure 4D). To directly assess the role of beauvericin in disease development, pot experiments were conducted in which beauvericin (1 μg g^−1^ soil) was exogenously applied to the soil. Beauvericin amendment significantly increased Fusarium wilt disease severity in tomato plants grown in natural soil (Welch’s *t*-test, *P* = 1.42 × 10^−3^, *P* = 3.69 × 10^−6^, and *P* = 7.53 × 10^−5^ for wild-type FOL, Cured-FOL, and Restored-FOL, respectively; Figure 4E). Notably, the disease-promoting effect of beauvericin was more pronounced in plants inoculated with Cured-FOL than in those inoculated with wild-type FOL or the Restored-FOL.

We next examined whether the bacterial endosymbiont EB05 directly produces beauvericin or instead promotes beauvericin biosynthesis by FOL. To test whether EB05 is capable of producing beauvericin, culture supernatants of EB05 were analyzed by HPLC-MS, and beauvericin was not detected (Figure S4A). Consistent with this result, PCR analysis detected the beauvericin synthase gene (*BEAS)* in the genomic DNA of wild-type FOL, Restored-FOL, and Cured-FOL, but not in EB05 (Figure S4B). In addition, genomic analysis of EB05 revealed no putative beauvericin biosynthetic gene cluster (Figure S4C). We then quantified *BEAS* gene expression in FOL strains. Expression of *BEAS* was higher in wild-type FOL and Restored-FOL than in Cured-FOL (Welch’s *t*-test, *P* = 1.19 × 10^−13^ and *P* = 6.62 × 10^−11^, respectively; Figure 4F), indicating that bacterial endosymbionts enhance beauvericin biosynthesis at the transcriptional level. To directly test the functional role of beauvericin in bacterial endosymbiont-mediated virulence, a *BEAS* deletion mutant was generated in the wild-type FOL background (designated as Δ*beas*-WT). Bacterial endosymbionts were subsequently removed from this mutant (Δ*beas*-Cured) and EB05 was reintroduced into the cured mutant (Δ*beas*-Restored). All these Δ*beas* strains lacked the capacity to produce beauvericin (Figure S4A) and caused lower Fusarium wilt disease severity compared with wild-type FOL and Restored-FOL (Tukey’s HSD test, *P* = 6.58 × 10^−15^; Figure S4D). Disease severity caused by Δ*beas*-Cured did not differ from that caused by Cured-FOL. Importantly, the magnitude of bacterial endosymbiont-mediated virulence significantly reduced in Δ*beas* mutants compared with FOL harboring intact *BEAS* gene (Tukey’s HSD test, *P* = 3.60 × 10^−8^; Figure 4G). In addition, soil microcosm and *in vitro* co-culture experiments showed that Δ*beas* mutants did not affect the abundance of St52 in soil and in oatmeal broth (Figure S4E). Consistent with these results, cell-free supernatants derived from Δ*beas* mutants did not inhibit St52 growth (Figure 4H).

### Beauvericin alters the rhizosphere microbiota and disrupts the disease-suppressiveness

We investigated the effects of exogenous beauvericin amendment on tomato rhizosphere bacterial communities (Figure 5A). In brief, amplicon sequencing revealed that beauvericin (1 μg g^−1^ soil) altered bacterial community β-diversity (PERMANOVA, *R*^2^ = 0.205, *P* = 8.30 × 10^−3^; Figure 5B). OTUs in this experiment were designated as beauvericin-responsive OTUs (be.OTUs). Beauvericin amendment resulted in significant increases and decreases in the relative abundances of 31 and 17 be.OTUs, respectively (Wald test, Benjamini-Hochberg adjusted *P* < 0.05; Figure S5). Notably, beauvericin treatment reduced relative abundance of four be.OTUs affiliated to *Streptomyces* sp. (*i*.*e*., be.OTU2496, be.OTU5527, be.OTU588, and be.OTU933) (Figure 5C). Comparison of these *Streptomyces* sp. be.OTUs with those identified in the FOL inoculation experiment (Figure 2C) showed that be.OTU2496, be.OTU5527, and be.OTU933 shared 100% partial 16S rRNA gene sequence identity with *Streptomyces* sp. f.OTU2295, f.OTU5685 and f.OTU273, respectively (Table S2). Consistent with these taxon-specific patterns, beauvericin amendment significantly reduced both the relative and absolute abundances of *Streptomyces* sp. in tomato rhizosphere (Welch’s *t*-test, *P* = 3.36 × 10^−5^ and *P* = 3.09 × 10^−5^, respectively; Figure 5D).

To evaluate the functional consequences of beauvericin-induced microbiota shifts, we assessed rhizosphere disease suppressiveness using a rhizosphere soil transplant experiment. Sterile field soil was amended with 3% (w/w) rhizosphere soil collected from beauvericin-treated or untreated pot experiments, after which a new generation of tomato plants was grown and challenged with wild-type FOL (Figure 5A). Tomato plants grown in soils containing beauvericin-treated rhizosphere inoculum exhibited significantly higher Fusarium wilt disease severity and increased FOL abundance in the rhizosphere compared to untreated rhizosphere inoculum (Welch’s *t*-test, *P* = 2.88 × 10^−4^ and *P* = 1.60 × 10^−3^, respectively; Figure 5E).

## DISCUSSION

Most eukaryotes form symbiotic relationships with diverse bacterial taxa, with consequences for host physiology, reproduction, and fitness.^7,13,14,33–35^ Among fungi, bacterial endosymbionts have been documented across diverse lineages, including several *Fusarium* spp., such as *F. solani, F. keratoplasticum*, and *F. fujikuroi*.^16,29,36^ In this study, we identified *Achromobacter* sp. as a prevalent bacterial endosymbiont residing within the hyphae of the soil-borne pathogen FOL. Although a representative isolate of this bacterial endosymbiont (EB05) was nonpathogenic to tomato plants when inoculated alone, its presence within FOL enhanced the pathogen’s virulence. Mechanistically, we demonstrate that this enhancement is mediated through bacterial endosymbiont-stimulated biosynthesis of beauvericin by FOL. Elevated beauvericin production, in turn, enabled FOL to suppress key disease-suppressive taxa within the rhizosphere microbiota, thereby weakening a critical microbial barrier to infection (Figure 6). Collectively, these findings establish bacterial endosymbionts as active contributors to fungal pathogenesis, revealing a mechanism by which bacterial endosymbiont-driven metabolic reprogramming of the fungal host amplifies virulence through disruption of plant-beneficial microbiota.

### Bacterial endosymbionts alter the effects of FOL on the rhizosphere microbiota

The rhizosphere microbiota constitutes a critical first line of defense against soil-borne pathogens and plays a central role in maintaining plant health.^2,18^ In this study, we found that the difference in virulence between bacterial endosymbiont-free and endosymbiont-harboring FOL strains was more pronounced when plants were grown in natural soil than in sterile soil. This context dependency indicates that the contribution of bacterial endosymbionts to FOL virulence is closely linked to interactions with the resident rhizosphere microbiota. Consistent with this interpretation, inoculation with bacterial endosymbiont-harboring FOL strains (i.e., wild-type FOL and Restored-FOL) resulted in rhizosphere bacterial community shifts that differ from those caused by Cured-FOL (bacterial endosymbiont-free FOL). Together, these results suggest that bacterial endosymbionts of FOL actively modulate rhizosphere bacterial communities during FOL invasion. Previous studies have shown that bacterial endosymbionts can enhance the virulence of fungal phytopathogens through direct mechanisms, such as promoting fungal growth or contributing to the biosynthesis of phytotoxic compounds.^7,15,16^ Our results support an additional, indirect mechanism by which specific bacterial endosymbionts amplify fungal virulence. Specifically, we show that specific bacterial endosymbiont taxa induce beauvericin biosynthesis in FOL, which in turn disrupts the resident rhizosphere microbiota and reduces its disease-suppressive capacity. This mechanism highlights the rhizosphere microbiota itself as a functional target of bacterial endosymbiont-mediated pathogenicity.

Plants dynamically interact with a diverse assemblage of pathogenic, commensal, and mutualistic microorganisms in the rhizosphere.^1,2,24^ Notably, we observed that bacterial endosymbiont-harboring FOL, but not endosymbiont-free FOL, caused a pronounced reduction in the abundance of *Streptomyces* sp. in tomato rhizosphere. Members of the genus *Streptomyces* are ubiquitous in soils and are widely recognized for their roles in biological control of phytopathogens, including FOL.^37,38^ These bacteria can protect plants through direct (*e*.*g*., antibiosis and competition for resources) as well as indirect mechanisms (*e*.*g*., induction of plant systemic resistance).^38^ The *Streptomyces* sp. St52 isolated in this study, a representative taxon suppressed by bacterial endosymbiont-harboring FOL, can inhibit FOL growth *in vitro* and reduce Fusarium wilt disease severity in tomato plants when inoculated into the soil. Importantly, we detected no direct antagonistic effect of bacterial endosymbiont EB05 on St52. Instead, EB05 altered the metabolic output of its fungal host, resulting in the suppression of St52. These findings indicate that EB05 facilitates FOL invasion indirectly, by enabling the pathogen to inhibit key disease-suppressive taxa within the rhizosphere microbiota rather than through direct bacterial-bacterial antagonism.

### Bacterial endosymbionts stimulate fungal beauvericin biosynthesis to disrupt the rhizobiome

Beauvericin is a cyclic hexadepsipeptide produced by several fungi, including species of *Fusarium, Beauveria*, and *Isaria* spp.^39^ It exhibits broad bioactivity, with documented toxicity toward microorganisms, plants, and mammalians,^32,40–42^ and has been implicated as a virulence factor in pathogenic *F. oxysporum*, particularly during early stages of host infection.^40^ Consistent with a role in pathogenicity, our metabolomic analyses showed that bacterial endosymbiont-free FOL (Cured-FOL) displayed a distinct extracellular metabolite profile compared with bacterial endosymbiont-harboring strains (wild-type FOL and Restored-FOL), including reduced beauvericin content. Importantly, multiple lines of evidence indicate that beauvericin is synthesized by the fungal host and transcriptionally upregulated in the presence of the bacterial endosymbiont. The bacterial endosymbiont EB05 lacked the beauvericin synthase gene *BEAS* gene and did not encode a recognizable *BEAS* gene cluster, and beauvericin was not detected in EB05 culture supernatants. In contrast, *BEAS* gene was present in all FOL backgrounds, and *BEAS* expression, as well as beauvericin accumulation, were elevated in bacterial endosymbiont-harboring strains relative to bacterial endosymbiont-free FOL. Moreover, genetic disruption of beauvericin biosynthesis substantially reduced disease severity and attenuated the bacterial endosymbiont-dependent virulence effect, providing direct evidence that beauvericin is a key mediator of bacterial endosymbiont-enhanced pathogenicity.

Beyond its effects on the host plant, beauvericin also emerged as a mechanism for targeting the microbial barrier to infection. Beauvericin inhibited the growth of the plant-beneficial *Streptomyces* St52, altered rhizosphere bacterial community structure, and reduced microbial-mediated disease suppressiveness in rhizosphere transplant assay. Together, these results support the idea that FOL can disrupt the microbial layer of plant defense by suppressing disease-suppressive bacteria taxa within the rhizosphere microbiota.^2,20,27^ In this context, endosymbiosis appears to provide a competitive advantage to FOL by amplifying beauvericin production, thereby weakening antagonistic bacterial populations that otherwise constrain pathogen establishment. This interpretation is consistent with previous studies showing that bacterial endosymbionts can alter fungal secondary metabolism and promote the production of virulence-associated metabolites.^15,16^ More broadly, our findings align with the well-established capacity of fungi to remodel their secondary metabolism in response to biotic and abiotic environmental stimuli,^43,44^ and with reports that of beauvericin production in *Fusarium* spp. is environmentally inducible.^41^ Future studies that identify the molecular mechanisms underlying bacterial endosymbionts modulate the metabolism of FOL can expand our view of the function of bacterial endosymbionts.

In summary, our results identify a mechanistic basis for bacterial endosymbiont-enhanced virulence in FOL. Specifically, the presence of *Achromobacter* sp. EB05 within fungal hyphae stimulates beauvericin biosynthesis by FOL, which suppresses specific disease-suppressive bacterial taxa (e.g., *Streptomyces* sp. St52) in the rhizosphere, thereby facilitating Fusarium wilt disease development. This work expands current models of fungal-bacterial symbiosis in pathogenesis by demonstrating how bacterial endosymbionts facilitate their fungal host to subvert rhizosphere-based defenses through targeted disruption of beneficial microbes. From an applied perspective, these findings suggest that strategies aimed at reinforcing disease-suppressive microbiota or disrupting key endosymbiotic interactions that potentiate toxin production may provide complementary approaches for managing soil-borne diseases while reducing reliance on conventional chemical control strategies.

## RESOURCE AVAILABILITY

### Lead contact

Additional information request should be directed to the lead contact, Xingang Zhou.

### Materials availability

Plant materials and bacterial strains used in this study can be made available upon request.

## Data and code availability

The sequencing data of the rhizosphere bacterial community were deposited in the Sequence Read Archive at NCBI with the accession number PRJNA1057741. The partial 16S rRNA gene sequences of isolated bacteria were submitted to NCBI GenBank with the accession numbers PP052935 to PP052952. The genome of *Achromobacter* sp. EB05 was deposited under BioProject accession CP155917. The R scripts utilized for statistical analyses and plotting figures are available at https://github.com/xingangzhouneau/endosymbiont.

## ACKNOWLEDGMENTS

This research was supported by the National Natural Science Foundation of China [42325704 (Z.W.), 32573013 (X. Zhou.)], the National Key Research and Development Program [2024YFD2300700 (X. Zhou.)], and the USDA National Institute of Food and Agriculture and Hatch Appropriations [Project PEN04908 and Accession 7006279 (F.D.-A.)].

## AUTHOR CONTRIBUTIONS

X. Zhou, Z.W., and F.W. conceived and designed the experiments; X. Zhou, X. Zhang, H.J., L.R., D.L., J.Z., and N.Z. performed the experiments and analyzed the data; X. Zhou, X. Zhang, M.K.u.R., Z.W., and F.D-A. wrote the manuscript. All authors edited the manuscript and approved the final version.

